# AMPBAN: A Deep Learning Framework Integrating Protein Sequence and Structural Features for Antimicrobial Peptide Prediction

**DOI:** 10.64898/2026.01.20.700468

**Authors:** Wenhui Bai, Wenhao Yang, Yao-Qing Chen, Hongchao Ji, Lorraine Brennan, Li Wang

**Affiliations:** Shenzhen Branch, Guangdong Laboratory of Lingnan Modern Agriculture, Agricultural Genomics Institute, Chinese Academy of Agricultural Sciences, Shenzhen, 518120, China; UCD Institute of Food and Health, UCD School of Agriculture and Food Science, University College Dublin, Dublin, Ireland; School of Public Health (Shenzhen), Shenzhen Campus of Sun Yat-sen University, Shenzhen, China

**Keywords:** antimicrobial peptides, machine learning, protein language model, equivariant graph neural network, bilinear attention network

## Abstract

The escalating crisis of antimicrobial resistance poses a devastating and immediate threat to human life. Antimicrobial peptides (AMPs) are a promising antibiotic substitute to combat antimicrobial resistance. Compared with the traditional wet-lab screening approaches, computational models have largely improved the efficiency of predicting antimicrobial peptide. However, most computational models overlook or underutilize the evolution and structural information of peptides, which is crucial for understanding the peptide functions. Here, we proposed a sophisticated deep learning model to predict AMPs, Antimicrobial Peptide Bilinear Attention Network (AMPBAN), which incorporates peptide evolution features from ESM3 protein language model, structure features from ESMFold predicted with equivariant graph neural network (EGNN), and the joint information from sequence and structure learned via Bilinear Attention Network. AMPBAN consistently demonstrated superior accuracy and generalization compared to nine state-of-the-art AMP prediction models across multiple independent benchmarks. Furthermore, an ablation study confirms that our multimodal fusion strategy significantly refines the integration of sequence and structural signals, yielding superior predictive balance over single-modality models. This framework provides a robust tool for the accelerated discovery of novel AMPs and the advancement of next-generation antimicrobial drug development. The datasets, source code and models are available at https://github.com/baiwenhuim/ampban.

## Introduction

The crisis of antimicrobial resistance (AMR) represents a major global health threat, responsible for an estimated 4.71 million deaths in 2021. Projections indicate it will remain a leading cause of mortality for decades [1]. This escalating resistance is compounded by a stagnation in the discovery and development of new antibiotics, creating a critical therapeutic gap [2–4]. To address this urgent need, Antimicrobial Peptides (AMPs), which are small, cationic, and amphipathic molecules, are gaining attention as promising alternatives [5]. AMPs are fundamental components of the innate immune system and exert broad-spectrum activity by physically interacting with microbial membranes, such as through pore formation and membrane destabilization, a mechanism that offers a lower likelihood of rapid resistance development compared to conventional, target-specific antibiotics [6,7]. Despite their significant therapeutic potential, as evidenced by over 85 peptide-based drugs approved by the United States Food and Drug Administration (USDA) and several AMPs currently undergoing clinical evaluation [8,9], the experimental discovery and validation of novel AMPs remain a costly, labor-intensive, and time-consuming process, highlighting the need for efficient computational tools to accelerate their identification.

To accelerate AMP discovery, early machine learning models leveraged manually engineered features combined with traditional algorithms like support vector machine (SVM), random forest (RF), and k-nearest neighbor (KNN) [10–12]. One key step of this classical approach is the development of handcrafted descriptors, often relying on Amino Acid Composition (AAC), Pseudo AAC (PseAAC), and key physicochemical properties. For instance, iAMP-2L employed PseAAC with fuzzy KNN [10], while AmPEP and MLAMP utilized the distribution patterns of amino acid properties and Grey PseAAC, respectively, in combination with RF [13,14]. Other models, such as iAMPpred and PTPAMP, further integrated compositional, physicochemical, and structural indices to train SVM [15,16]. While effective, the performance of these methods relies heavily on the quality and suitability of the chosen features, often leading to a tendency for feature redundancy or insufficient information to capture the complexity of AMP function, thereby limiting generalization and sometimes resulting in high false-positive rates [17].

The advent of deep learning methods has profoundly advanced AMP prediction by enabling models to learn hierarchical representations directly from peptide sequences. These methods aim to extract global features from peptides using advanced deep neural networks and other related algorithms, including convolutional neural networks (CNNs), recurrent neural networks (RNNs), long short-term memory (LSTM) models, and attention-based networks [18–20]. For example, Deep-AmPEP30 used a CNN on composition-transition-distribution features [18], while AMPlify and AMP-CLIP leveraged Bi-LSTMs and attention mechanisms to extract rich sequence embeddings [20,21]. More advanced sequence-centric models like ACEP [22] and iAMP-Attenpred [23] incorporated position-specific scoring matrices (PSSM) or pre-trained BERT embeddings into complex hybrid architectures involving CNNs, LSTMs, and attention. Despite their success, these sequence-centric models are inherently limited in their ability to capture the crucial structural determinants of antimicrobial activity, which are essential for modulating a peptide’s function [24].

Recent breakthroughs in protein structure prediction and graph neural networks (GNNs) offer a promising opportunity to bridge this gap. Tools like AlphaFold and ESMFold can generate highly accurate 3D protein and peptide structures, while GNNs are well-suited for encoding the spatial and relational information present in these structures [25–27]. Several pioneering studies have attempted to integrate structural information into AMP prediction. For instance, sAMPpred-GAT and similar models have used GNNs to leverage predicted protein contact maps, which represent inter-residue distances [28]. A notable recent example is the SSFGM-Model [27], which combined Molecular Surface Interaction Fingerprinting (MaSIF) features extracted from AlphaFold2-predicted peptide structures with sequence-based features into a GNN and geometric neural network to identify functional AMPs. Concurrently, protein language models (PLMs) such as ESM and ProtBERT have demonstrated remarkable capability in capturing contextual and evolutionary information from vast protein sequence datasets [29–31]. However, a significant limitation persists: most models that attempt to combine these powerful modalities, such as deepAMPNet [32] and TP-LMMSG [33], utilize only a limited subset of structural data, such as *C_α_ - C_α_* or *C_β_ - C_β_* distances. This approach strictly overlooks other rich structural cues, including complete 3D coordinate information and the intricate spatial relationships between sequence and structure, which are critical for comprehensive functional analysis.

In this work, we propose AMPBAN (Antimicrobial Peptide Bilinear Attention Network), a deep learning framework that integrates both sequence and structural features for robust AMP prediction. AMPBAN combines contextual sequence embeddings from the state-of-the-art ESM3 protein language model with 3D structural representations predicted by ESMFold and encoded using equivariant graph neural network (EGNN). To effectively model the interactions between these modalities, we introduce a bilinear attention network (BAN) that learns joint representations of sequence–structure interactions. By integrating evolutionary, structural, and relational information, AMPBAN provides a more holistic representation of peptides, enabling accurate and generalizable AMP discovery.

## MATERIALS AND METHODS

### 2.1 Framework of AMPBAN

In this study, we employed a Bilinear Attention Network (BAN) to integrate sequence and 3D structural features to enhance AMP prediction, addressing the limitations of methods that overlook structural details, and facilitating informative peptide sequence and structure embedding. Initially, peptide sequences are embedded using ESM3, producing 1920-dimensional features, which represent evolutionary information. For structural feature extraction, 3D protein structures generated by ESMFold are processed using E(n)-Equivariant Graph Neural Network (EGNN), which maintain equivariance to translations and rotations, ensuring accurate geometric modeling while capturing peptide structures with fine granularity. A Bilinear Attention Network then fuses sequence and structural data by computing pairwise attention weights between their feature channels, followed by bilinear pooling to create a joint feature vector. Subsequently, the sequence feature vector, structural feature vector from EGNN, and joint feature vector from BAN are concatenated. This composite feature vector is then fed into a fully connected neural network, which outputs the probability of the peptide being an AMP through a sigmoid activation function. AMPBAN facilitates precise modeling of peptide structures at a high resolution, enabling the differentiation of amino acids based on their unique characteristics, variations, and roles in peptide attributes, such as antimicrobial function. The model was subsequently trained on annotated datasets to execute a binary classification task. (Figure. 1). The training parameters for AMPBAN are comprehensively detailed in Supplementary Text S1.

**Figure 1.**
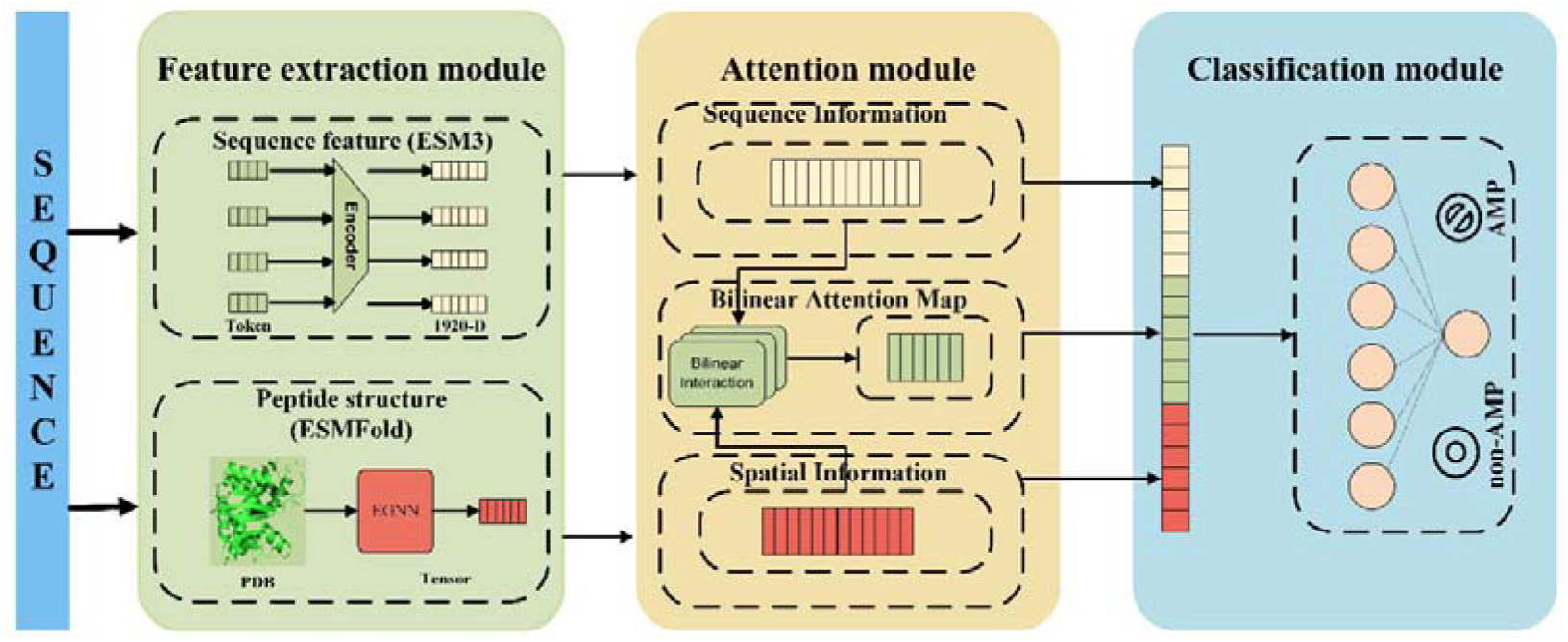
Overview of the AMPBAN framework. **Feature extraction module:** Peptide sequences are processed by ESM3 to extract sequence-based features. Concurrently, the 3D protein structure is generated from the sequence via ESMFold. The structural data is then processed by an E(n)-Equivariant Graph Neural Network (EGNN) to derive structural features. **Attention module:** A Bilinear Attention Network (BAN) integrates the sequence features from ESM3 with the structural features from the EGNN. A bilinear pooling operation follows, producing a joint feature vector that captures the interaction between sequence and structural information. **Classification module:** The final composite feature vector is formed by concatenating the sequence feature vector (from ESM3), the structural feature vector (from EGNN), and the joint feature vector (from BAN). This comprehensive vector is put into a fully connected neural network, which performs binary classification to predict the probability of a peptide being an antimicrobial peptide or not.

### 2.2 Benchmark datasets

To establish a robust training dataset, we curated the up-to-date antimicrobial peptide data set from Peptipedia v2.0 [34], a widely recognized and comprehensive public repository of peptides which include 76 AMP source from public database, like Swiss-Prot [35], LAMP2 [36], DBAASP [37], PeptideDB [38], CAMP [39], DRAMP [40], PlantPepDB [41], APD3 [7] e.g., to amount datasets of training AMP prediction model, like Ampfun [42], AMPlify [20], AMP Scanner 2 [19], APIN [43] e.g. From this extensive resource, we identified an initial set of 103,561 peptides with documented biological activity. Following a stringent filtering process to retain only sequences composed of canonical amino acids, we obtained a refined dataset comprising 32,846 AMP sequences, with lengths ranging from 5 to 100 amino acid residues.

For the generation of non-AMP sequences, we retrieved 58,095 manually annotated sequences from the UniProtKB/Swiss-Prot database (2024_12 release) [44]. To ensure a high degree of confidence in their non-antimicrobial nature, we applied rigorous filtering criteria, resulting in a final set of 42,163 non-AMP sequences. The full description of the filtering criteria is provided in the Supplementary Text S2.

Crucially, to prevent any potential data leakage between the training data and the three independent test datasets, we implemented a filtering step using a Python script. This script identified and removed any sequences present in the training data that were also found in any of the independent test sets. Following this rigorous data cleaning procedure, the final training dataset consisted of 30,971 AMP sequences and 38,675 non-AMP sequences (Table 1).

**Table 1.**
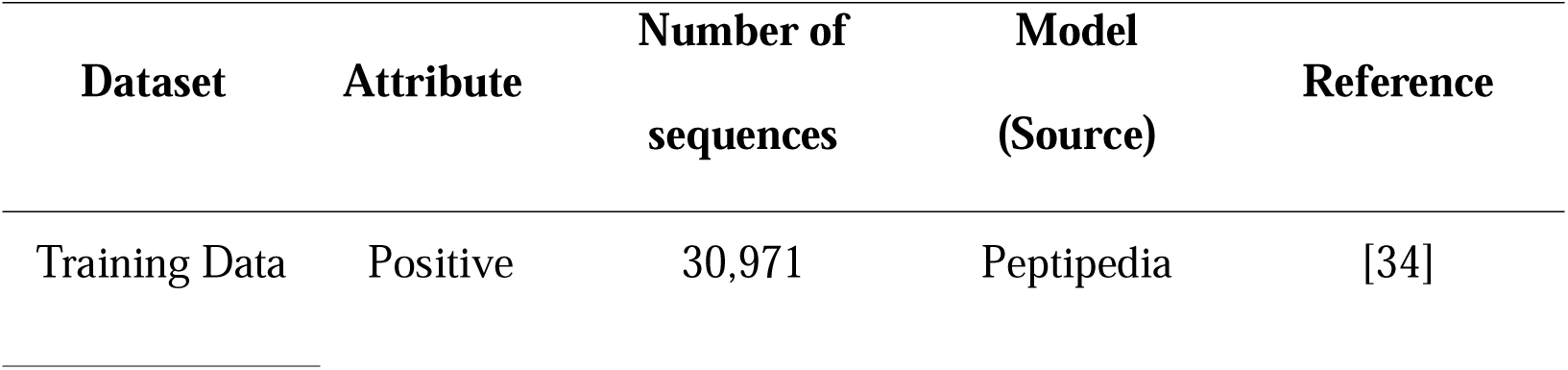
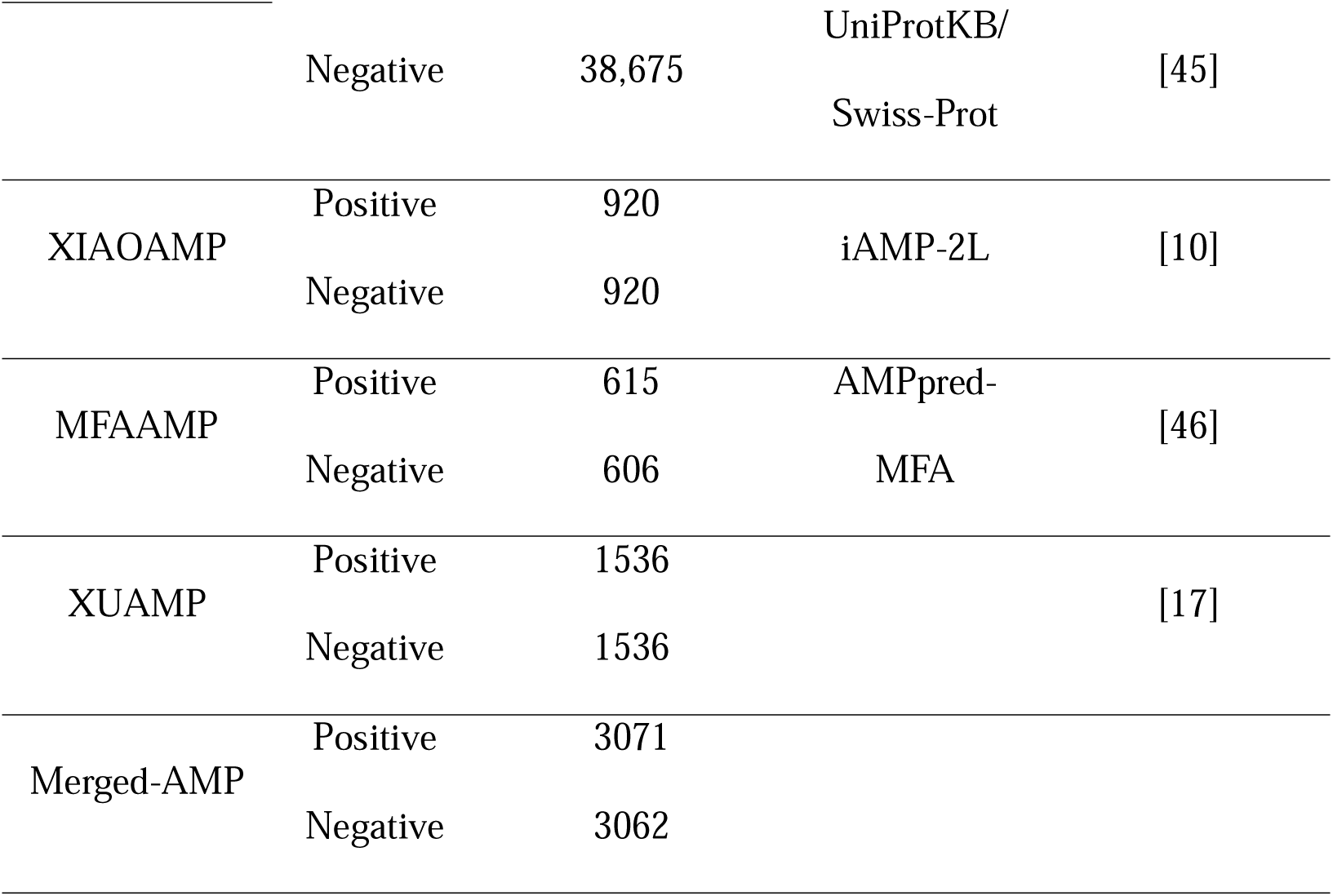
The characteristics of both the training and test dataset.

### 2.3 Independent test set

To validate the objective robustness and generalization capability of the proposed model in identifying novel sequences, we constructed a comprehensive independent test set designated as Merged-AMP. This dataset consolidates three balanced independent test sets that have been utilized by more than ten existing prediction methods, ensuring an unbiased and equitable benchmark for performance comparison. The first component, designated as XIAOAMP (originally employed by Xiao et al. [10]), consists of 920 AMP sequences sourced from the Antimicrobial Peptide Database (APD3) [7] and 920 non-AMP sequences known to lack antimicrobial activity. The second dataset, termed MFAAMP (originally used in the development of AMPpred-MFA [46]), comprises 615 AMP sequences and 606 non-AMP sequences.

The third dataset, referred to as XUAMP (reported by Xu et al. [17]), includes 1,536 AMP sequences and 1,536 non-AMP sequences (Table 1).

The peptide sequences in this merged dataset range from 5 to 102 residues in length, with an average length of 47 and a mode of 33. Additionally, we analyzed the length distribution and found that these AMPs are evenly distributed across each length range (Figure S1).

### 2.4 Sequence Feature Extraction: Evolutionary Scale Modeling

For each peptide sequence, we employed ESM3 (also referred to as ESMC), a state-of-the-art Transformer-based protein language model developed by Meta AI, to generate sequence embeddings [29]. ESM3 is pre-trained on the UniRef50 dataset [45] using a masked language modeling approach, enabling it to capture intricate contextual relationships and deep evolutionary information within amino acid sequences.

To obtain the final fixed-length feature vector (*X_seq_*) used for prediction and fusion, we utilized ESM++ (a Hugging face compatible implementation of ESM3) with the small model (300 million parameters), which produces residue-level embeddings of 960 dimensions. For a sequence of length *L*, the output is a matrix *X*_residue_ *E R*^(Lx960)^. We derive the sequence-level features by applying two distinct pooling methods to the residue embeddings: Mean Pooling, which averages the embedding across all *L* residues to yield a 960-D vector, and [CLS] Pooling, which leverages the dedicated [CLS] token embedding to generate another 960-D vector. These two 960-D vectors are concatenated to form the final 1920-dimensional feature vector (*X*_seq_). This vector serves as the sequence input to the BAN and is also one of the components concatenated for the final classifier.

### 2.5 Structure Feature Extraction: Equivariant Graph Neural Networks

To incorporate 3D structural information into the AMPBAN model, we first predicted the tertiary structures of all candidates’ AMPs using ESMFold, a state-of-the-art protein structure prediction tool developed by Meta AI. ESMFold leverages the ESM language model to generate high-quality protein structures from sequence alone, outperforming traditional methods like AlphaFold2 in speed while maintaining comparable accuracy for short sequences like AMPs. For each peptide sequence in our dataset, we input the amino acid sequence into ESMFold v1.0, which outputs a predicted 3D coordinate file in PDB format. The prediction process was run on a GPU-enabled system to handle the computational load, with default parameters (No template usage, as AMPs are typically short and do not require homology modeling). This step resulted in predicted structures for all training (positive and negative AMPs) and independent test sequences, enabling the subsequent extraction of geometry-aware features.

Following structure prediction, we employed the Progres package (version 1.0) to generate low-dimensional structure graph embeddings for each peptide. Progres, developed by Greener and Jamali, is a fast protein structure searching tool that uses an E(n)-equivariant graph neural network (EGNN) to embed protein domains into a 128-dimensional representation, facilitating rapid structural comparisons independent of primary sequence. EGNNs are a class of graph neural networks that are equivariant to rotations, translations, reflections, and permutations, as introduced by [47]. They extend standard GNNs by incorporating coordinate updates alongside feature embeddings, ensuring that transformations on input coordinates result in equivalent transformations on outputs.

The EGNN in Progres treats protein structures as graphs where *C_α_* atoms serve as nodes, with edges connecting *C_α_* atoms within a 10 Å distance threshold. Each node incorporates the following features: (1) the number of *C_α_* atoms within 10 Å, normalized by the maximum such number in the protein; (2) indicators for N-terminal or C-terminal positions; (3) the *r* torsion angle between consecutive *C_α_* atoms; and (4) a 64-dimensional sinusoidal positional encoding based on residue number. The model consists of six Equivariant Graph Convolutional Layers (EGCLs) with residual connections, preceded by a one-layer multilayer perception (MLP) on node features and followed by a two-layer MLP. Each hidden layer has 128 dimensions and uses the Swish/SiLU activation function, except for the edge MLP in the EGNN, which has a hidden layer of 256 dimensions and a 64-dimensional output. Node features are sum-pooled, and a final two-layer MLP produces 128-dimensional embedding, ensuring SE(3)-invariance (no change with translation or rotation).

The EGCL operates as follows: For each edge *(i, j)*, the edge embedding *m*_ij_ is computed as

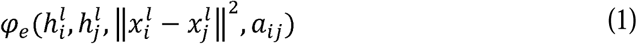

where 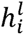 is the node feature embedding at layer l, 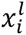 is the coordinate embedding *α_ij_* is the edge attribute, and *φ_e_* is an MLP. The coordinate update is

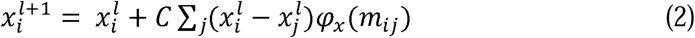

where *φ_x_* is an MLP and C is a normalization factor (*1/(M - 1)* for *M* nodes). The node embedding update is

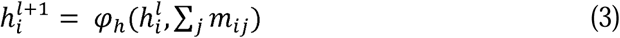

where *φ_h_* is an MLP. This design preserves E(n) equivariance: translations and rotations on inputs result in equivalent transformations on outputs, while permutations of nodes are handled through message passing. The 128-dimensional embedding (*X_struct_*) captures geometric and topological properties, enabling the AMPBAN model to fuse structural information with sequence features for improved prediction accuracy. The Progres embeddings were normalized before integration into the BAN layer.

### 2.6 Bilinear Attention Network

To effectively leverage both the high-dimensional peptide sequence embedding (*X_seq_*) and the graph structure feature (*X_struct_*) for AMP prediction, we employ the Bilinear Attention Network (BAN).

Our implementation of BAN utilizes Multi-Modal Low-Rank Bilinear Pooling (MRL-B), which allows for the efficient calculation of the outer product interactions between the two input modalities, *X_seq_* (Query) and *X_struct_* (Value). This strategy drastically reduces computational complexity while retaining the expressive power of full bilinear pooling.

#### Low-Rank Projection and Interaction Map

Instead of directly computing a full bilinear attention map, the BAN module first projects the two input feature vectors into a shared, lower-dimensional space using *k* parallel linear sub-networks (where *k* is typically set to 3).

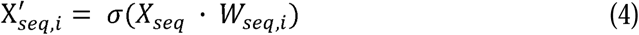

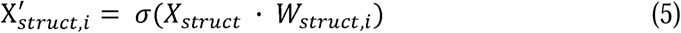

where *W_seq,i_* and *W_struct,i_* are the weight matrices for the *i*-th sub-network, and *a* is the ReLU activation. This process yields *k* pairs of projected vectors.

The bilinear interaction map *A* is implicitly calculated by combining these low-rank projections: the attention strength between the two global features is derived from the outer product of their projected forms. This interaction map effectively computes the element-wise similarity and synergy between the corresponding dimensions of the sequence and structural features, highlighting which combination of feature channels is most predictive of antimicrobial function.

#### Bilinear Pooling and Dimensionality Reduction

The bilinear pooling layer then aggregates the information captured in the interaction map *A* across all *k* sub-networks to produce the compact joint feature vector, *f_joint_*. The core feature fusion is accomplished by taking the inner product of the projected sequence and structure vectors weighted by the calculated attention map. Since *k* sub-networks are used, the resulting vector *f*, has a total dimension of *k* × *h_dim_*. A Sum Pooling operation is performed over this vector with a stride equal to *k*.

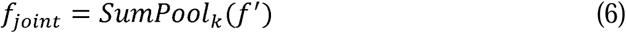

This pooling step reduces the dimensionality back down to *h_dim_* (the hidden dimension), resulting in a single, fixed-size vector *f_joint_* that comprehensively encodes the multimodal relationship between the peptide’s sequence and 3D structure. The feature is then processed by a Batch Normalization layer before being passed to the final classifier.

### 2.7 Classification module

Following the feature fusion via BAN, the final classification module integrates three distinct feature modalities for AMP prediction. The input to the classifier is a concatenated composite vector, aggregating the 1920-dimensional sequence embedding (*X*_seq_) from ESM3, the 128-dimensional structural embedding (*X_struct_*) from EGNN, and the 128-dimensional joint feature vector (*f_joint_*) output by the BAN layer. This 2176-dimensional vector is first mapped to a 128-dimensional hidden space, followed by a Batch Normalization layer to standardize feature distribution, accelerate convergence, mitigate gradient vanishing, and enhance model stability. A ReLU activation function is then applied to introduce nonlinearity, enabling the capture of complex relationships between fused features and antimicrobial activity. A Dropout layer is incorporated post-ReLU to prevent overfitting, followed by a linear layer that reduces the hidden representation to a single logit. Finally, a Sigmoid activation function transforms the logit into a probability score between 0 and 1, indicating the likelihood of the peptide being an AMP.

### 2.8 Performance Evaluation Metrics

To evaluate the performance of the proposed model in antimicrobial peptide (AMP) recognition, several metrics, including accuracy, sensitivity, specificity, Matthew’s correlation coefficient (MCC), precision, area under the curve (AUC) were used to assess both the proposed model and baseline methods. The formulas for these metrics are as follows:

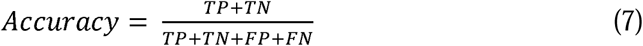

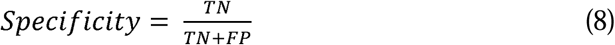

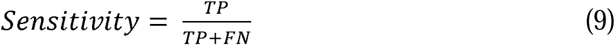

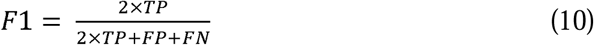

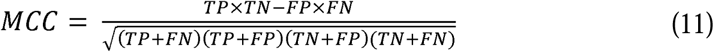

where True Positive (TP) refers to the number of samples that are positive and are predicted as positive by the model. True Negative (TN) refers to the number of samples that are negative and are predicted as negative by the model. False Positive (FP) refers to the number of samples that are negative but are predicted as positive by the model. False Negative (FN) refers to the number of samples that are positive but are predicted as negative by the model.

## 3 RESULTS AND DISCUSSION

### 3.1 Sequence Analysis of Collected AMPs

A total of 30,971 AMPs and 38,675 non-AMPs were used to construct the AMP classifier. We first compared the amino acid compositions and physicochemical properties between AMPs and non-AMPs. We found significant differences in the amino acid frequency between the AMPs and non-AMPs (Figure 1A). AMPs usually contain a higher proportion of positively charged amino acids, such as arginine (R) and lysine (K), hydrophobic isoleucine (I), stable glycine (G) and cysteine (C), which facilitate the formation of a stable state and interact with the negatively charged phospholipid head groups on the bacterial cell membrane, thereby destroying the integrity of the cell membrane [49]. This amino acid composition is consistent with what was previously reported [42,46]. Among the surveyed physicochemical properties, sequence length, net charge, hydropathicity, isoelectric point, molecular weight, instability index, and the secondary structure fraction (helix and sheet) exhibited profoundly significant differences (p ≤ p ≤ 0.0001) (Figure 1B-E; Figure S2; Supplementary Text S3). AMP sequences are significantly shorter than non-AMP sequences and exhibit a narrow range of lengths, while non-AMP sequences have a wide distributional range of lengths.

Altogether, these findings highlight the unique and distinct structural and compositional features of AMPs compared with non-AMPs (Figure 2), which may be related to their biological functions and mechanisms of action.

**Figure 2.**
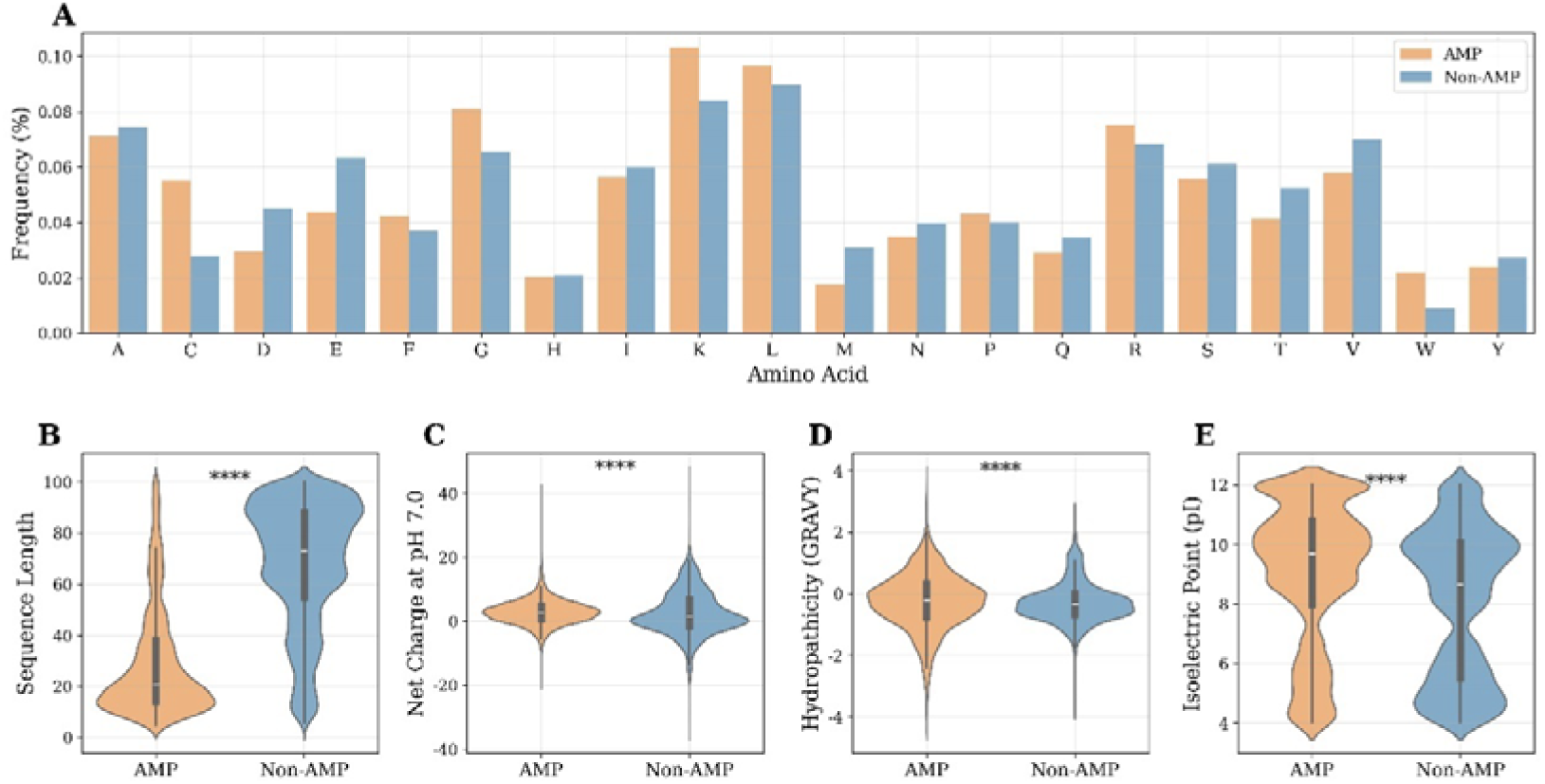
Physicochemical properties and amino acid composition of AMPs and Non-AMPs. (A) Amino Acid Frequency: A bar chart illustrating the percentage frequency of each amino acid (A, C, D, etc.) for both AMPs (Orange bars) and non-AMPs (blue bars). (B-E) Physicochemical Property Distributions: Violin plots showing the statistical distribution of four key physicochemical properties (Sequence Length, Net Charge at PH=7.0, Hydropathicity (GRAVY Score), Isoelectric Point) for both AMPs and non-AMPs. Each plot displays density distribution, with the white dot representing the median and the thick black bar indicating the interquartile range.

### 3.2 Similarity Between the Benchmark data and Independent Test datasets

To evaluate the consistency between the training data and three external test datasets (MFAAMP, XUAMP, and XIAOAMP) [10,17,46], we conducted a comprehensive comparison of physicochemical and compositional features, including eight key physicochemical properties and the fractional composition of all 20 canonical amino acids. Uniform Manifold Approximation and Projection (UMAP) reveals substantial overlap between the training and merged independent data (Figure 3A), indicating strong similarities and a comprehensive representation of the training data. Notably, the XUAMP and MFAAMP (Figure 3B and 3D) exhibit close alignment with the training manifold, while the XIAOAMP (Figure 3C) displays moderate separation, particularly in regions enriched for longer, charged peptides. The observed divergence with the XIAOAMP suggests potential domain-specific biases (such as sequence length or charge extremes), warranting cautious interpretation of performance on this cohort.

**Figure 3.**
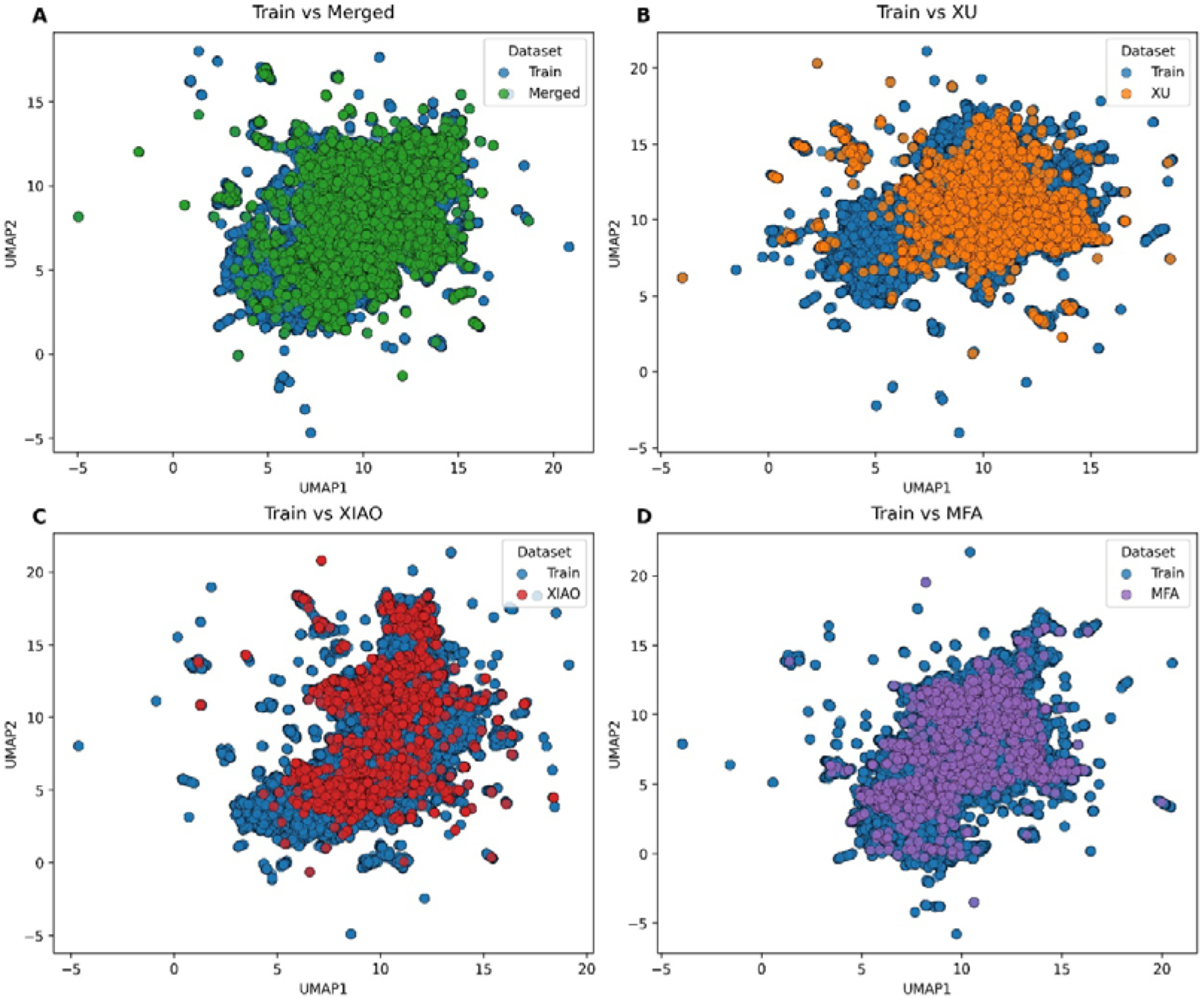
The similarity between training datasets and three independent test datasets (MFAAMP, XUAMP, and XIAOAMP). (A) Training set vs. merged independent test sets (MFAAMP + XUAMP + XIAOAMP). (B) Training set vs. XUAMP. (C) Training set vs. XIAOAMP. (D) Training set vs. MFAAMP.

These findings confirm that the training corpus encompasses the major of physicochemical and sequence compositional patterns present in the independent benchmarks. The distributional congruence demonstrates compelling evidence of the use of the training set as a robust foundation for AMP prediction across diverse external evaluations, implying the potential generalizability of models trained on this dataset.

### 3.3 Performance Evaluation Based on the Benchmark Data Sets

To rigorously validate the effectiveness and superiority of the proposed AMPBAN model, and ensure an unbiased evaluation, we meticulously re-implemented four advanced AMP prediction models—AMPScanner, AMPlify, Macrel, and AMP-CLIP—by strictly adhering to the original data processing protocols, feature extraction strategies, and model architectures described in the respective source papers [19–21,50]. The detailed implementation procedures were elaborated in Supplementary Text S4. All models, including AMPBAN, were trained and evaluated with identical 5-fold cross-validation on our curated benchmark dataset, eliminating potential biases arising from data splitting, preprocessing discrepancies, or hyperparameter inconsistencies. The mean and standard deviation of six comprehensive metrics, including Accuracy (ACC), Matthews Correlation Coefficient (MCC), Sensitivity (Sn), Specificity (Sp), F1-score, and Area Under the ROC Curve (AUC), are compared.

AMPBAN consistently outperformed all four baselines across most key metrics, achieving the highest Accuracy (0.9302 ± 0.0016), MCC (0.8606 ± 0.0038), Sensitivity (0.9463 ± 0.0060), F1-score (0.9236 ± 0.0020), and AUC (0.9785 ± 0.0009) (Table 2). While AMPBAN’s Specificity was 0.9177 ± 0.0043, it was marginally surpassed by AMPScanner (0.9250 ± 0.0061) and AMPlify (0.9221 ± 0.0090), but its substantially higher Sensitivity confirms its stronger ability to detect true positives (Table 2). AMPlify ranked second with competitive performance (Acc: 0.9261 ± 0.0016, AUC: 0.9752 ± 0.0010), yet surpassed by AMPBAN in Sensitivity and MCC, highlighting AMPBAN’s superior predictive balance. Macrel and AMPScanner delivered solid results but lagged 0.02–0.03 behind AMPBAN in key indicators (Table 2), reflecting limitations of manual feature engineering and hybrid convolutional-recurrent designs on complex sequence patterns. Notably, AMP-CLIP exhibited the poorest and most unstable performance (Sensitivity: 0.3306 ± 0.4619, MCC: 0.1974 ± 0.2794), likely due to its original design for contrastive pre-training rather than direct binary classification. These reproducible results conclusively demonstrate that AMPBAN, without relying on extensive handcrafted features or large-scale pre-training, establishes a new state-of-the-art benchmark for AMP prediction, offering an accurate, robust, and practical tool for large-scale AMP discovery.

**Table 2.** Performance comparison of AMPBAN with reproduced methods on the Benchmark dataset. The bold values showed the best performance.

| Model | ACC | AUC | F1 | MCC | SN | SP |
| --- | --- | --- | --- | --- | --- | --- |
| AMPScan | 0.9200 ± | 0.9704 ± | 0.9066 ± | 0.8368 ± | 0.9133 ± | **0.9250 ±** |
| ner | 0.0012 | 0.0005 | 0.0017 | 0.0026 | 0.0084 | **0.0061** |
| AMP- | 0.6198 ± | 0.8504 ± | 0.2909 ± | 0.1974 ± | 0.3306 ± | 0.8515 ± |
| CLIP | 0.1060 | 0.0136 | 0.3989 | 0.2794 | 0.4619 | 0.2710 |
| Macrel | 0.9071 ± | 0.9657 ± | 0.8983 ± | 0.8128 ± | 0.8982 ± | 0.9146 ± |
|  | 0.0033 | 0.0024 | 0.0038 | 0.0067 | 0.0064 | 0.0027 |
| AMPlify | 0.9261 ± | 0.9752 ± | 0.9180 ± | 0.8512 ± | 0.9310 ± | 0.9221 ± |
|  | 0.0016 | 0.0010 | 0.0026 | 0.0036 | 0.0135 | 0.0090 |
| AMPBAN | **0.9302 ±** | **0.9785 ±** | **0.9236 ±** | **0.8606 ±** | **0.9463 ±** | 0.9177 ± |
|  | **0.0016** | **0.0009** | **0.0020** | **0.0038** | **0.0060** | 0.0043 |

### 3.4 Performance Evaluation Based on the Independent Test Data Sets

As AMPBAN showed superiority over four rigorously reproduced state-of-the-art models on the benchmark dataset through identical preprocessing, feature extraction, and 5-fold cross-validation protocols (Table 2), we next evaluated its generalization and robustness on independent test datasets, a more challenging and realistic scenario. Here we compare the performance of AMPBAN with nine advanced predictors, including five deep learning–based approaches CAMP3-ANN [11], AI4AMP [51], AMP-CLIP [21], AMPlify [20], and AMPScannerV2 [19], and four traditional machine learning methods AMPfun [42], smAMPsTK [52], PTPAMP[16], and MACREL [50]. All competing tools are either accessible through official web servers or reproducible via published source code (Table S1), ensuring fair and verifiable comparisons. Performance was assessed on three widely accepted independent test sets— XUAMP [17], XIAOAMP [10] and MFAAMP [32,46], as well as on a merged dataset combining all three.

On the merged independent test dataset, AMPBAN achieved the highest accuracy (0.8358), AUC (0.8897), MCC (0.6887), and specificity (0.9461), while securing the second-highest F1-score (0.8157) and sensitivity (0.7258) (Table 3 and Figure 4A). This superior balanced performance across diverse data sources underscores AMPBAN’s ability to effectively integrate sequence and structural information for robust AMP prediction.

**Figure 4.**
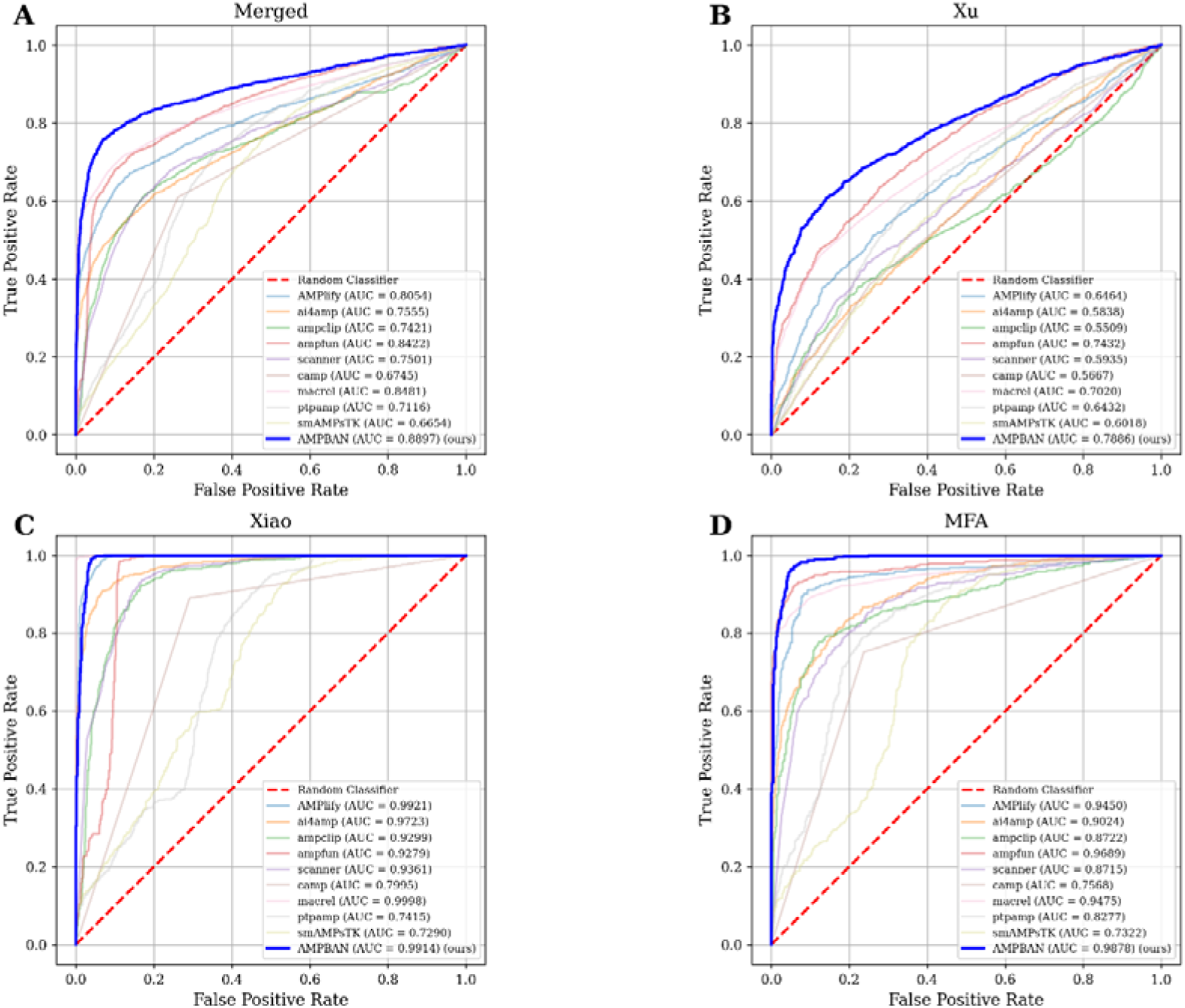
Receiver operating characteristic (ROC) curves of AMPBAN and the other nine predictors on the four independent test datasets. (A) Merged-AMP: A combined dataset consisting of the XUAMP, XIAOAMP, and MFAAMP. (B) XUAMP. (C) XIAOAMP. (D) MFAAMP. The dotted red line represents the performance of a Random Classifier (AUC = 0.5); the blue line indicates that of AMPBAN.

**Table 3.**
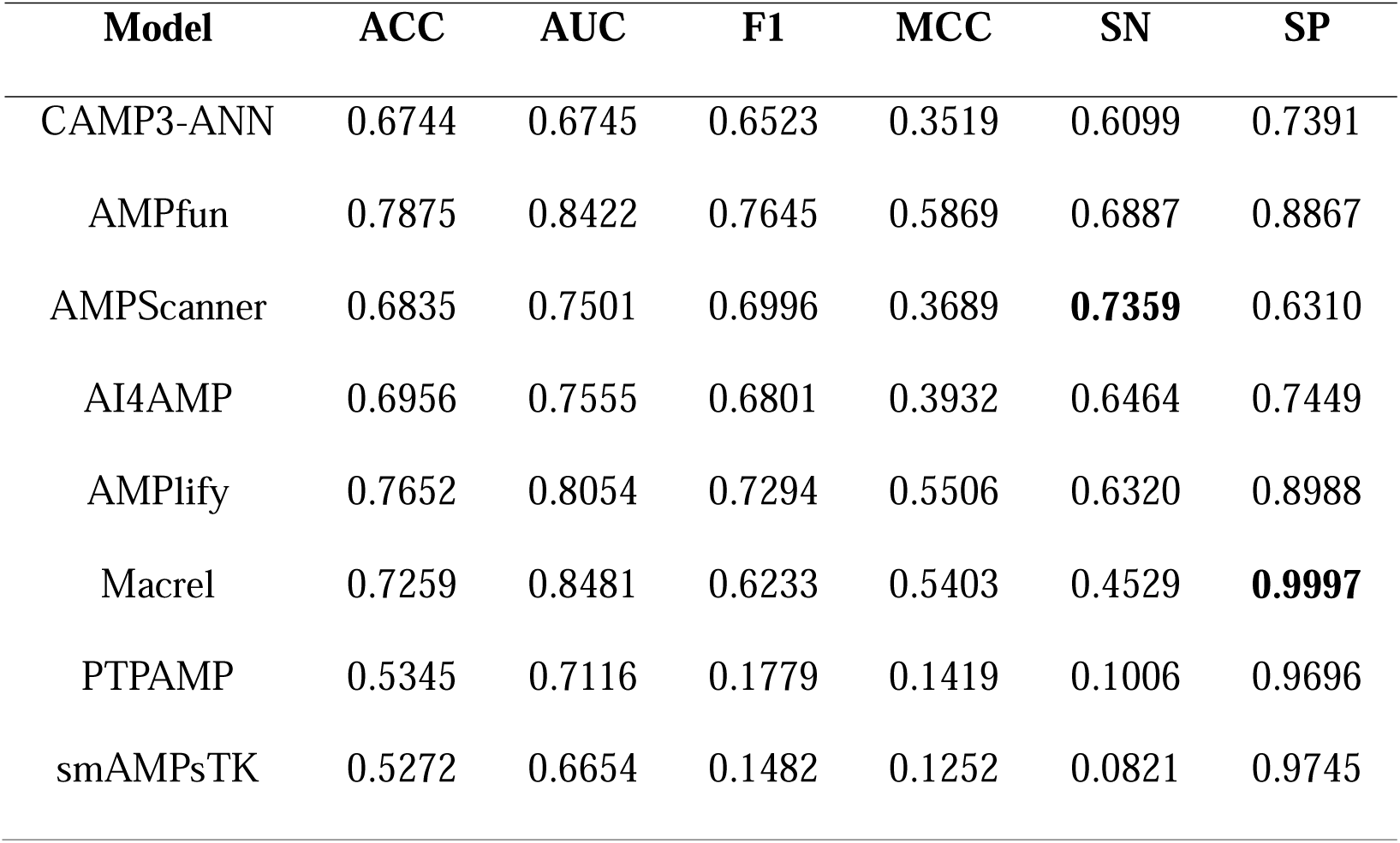
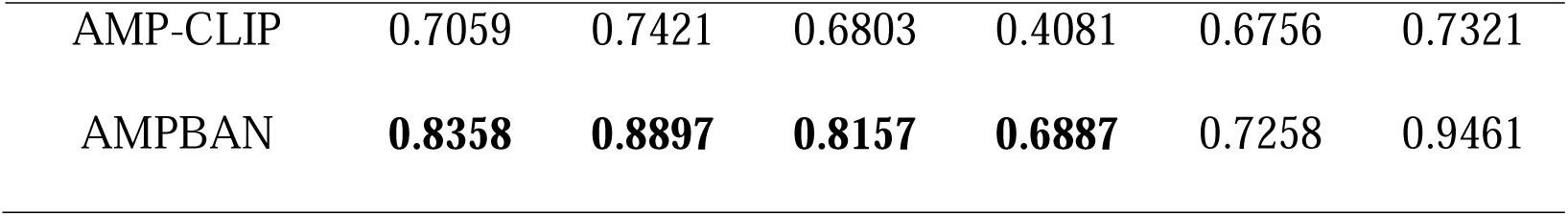
Performance of AMPBAN compared with existing methods on the Merged-AMP dataset. The bold values showed the best performance.

| Model | ACC | AUC | F1 | MCC | SN | SP |
| --- | --- | --- | --- | --- | --- | --- |
| CAMP3-ANN | 0.6744 | 0.6745 | 0.6523 | 0.3519 | 0.6099 | 0.7391 |
| AMPfun | 0.7875 | 0.8422 | 0.7645 | 0.5869 | 0.6887 | 0.8867 |
| AMPScanner | 0.6835 | 0.7501 | 0.6996 | 0.3689 | **0.7359** | 0.6310 |
| AI4AMP | 0.6956 | 0.7555 | 0.6801 | 0.3932 | 0.6464 | 0.7449 |
| AMPlify | 0.7652 | 0.8054 | 0.7294 | 0.5506 | 0.6320 | 0.8988 |
| Macrel | 0.7259 | 0.8481 | 0.6233 | 0.5403 | 0.4529 | **0.9997** |
| PTPAMP | 0.5345 | 0.7116 | 0.1779 | 0.1419 | 0.1006 | 0.9696 |
| smAMPsTK | 0.5272 | 0.6654 | 0.1482 | 0.1252 | 0.0821 | 0.9745 |
| AMP-CLIP | 0.7059 | 0.7421 | 0.6803 | 0.4081 | 0.6756 | 0.7321 |
| AMPBAN | **0.8358** | **0.8897** | **0.8157** | **0.6887** | 0.7258 | 0.9461 |

On the more challenging XUAMP, AMPBAN outperformed all competitors, attaining the highest accuracy (0.7077), MCC (0.4691), F1-score (0.6192), and specificity (0.9401) (Table 4 and Figure 4B), despite the dataset’s inherent difficulty. Although overall scores were lower than on XIAOAMP and MFAAMP, AMPBAN consistently surpassed both deep learning and traditional methods, demonstrating strong generalization under stringent conditions.

**Table 4.** Performance of AMPBAN compared with existing methods on XUAMP dataset. The bold values showed the best performance.

| Model | ACC | AUC | F1 | MCC | SN | SP |
| --- | --- | --- | --- | --- | --- | --- |
| CAMP3-ANN | 0.5667 | 0.5667 | 0.4708 | 0.1432 | 0.3854 | 0.7480 |
| AMPfun | 0.6566 | 0.7432 | 0.5415 | 0.3621 | 0.4056 | 0.9076 |
| AMPScanner | 0.5755 | 0.5935 | 0.5519 | 0.1519 | 0.5228 | 0.6283 |
| AI4AMP | 0.5566 | 0.5838 | 0.4543 | 0.1222 | 0.3691 | 0.7441 |
| AMPlify | 0.6032 | 0.6464 | 0.4364 | 0.2560 | 0.3073 | 0.8991 |
| Macrel | 0.5267 | 0.7020 | 0.1025 | 0.1637 | 0.0540 | 0.9993 |
| PTPAMP | 0.5137 | 0.6432 | 0.1118 | 0.0643 | 0.0612 | 0.9661 |
| smAMPsTK | 0.5052 | 0.6018 | 0.0743 | 0.0285 | 0.0397 | **0.9707** |
| AMP-CLIP | 0.5645 | 0.5509 | 0.4962 | 0.1339 | 0.4290 | 0.6999 |
| AMPBAN | **0.7077** | **0.7886** | **0.6192** | **0.4691** | **0.4753** | 0.9401 |

On the XIAOAMP, AMPBAN demonstrated competitive top-tier performance, achieving high accuracy (0.9723), MCC (0.9455), and F1-score (0.9729), alongside a robust specificity of 0.9500 (Table 5 and Figure 4C). Notably, AMPBAN maintained near-perfect sensitivity (0.9946), outperforming Macrel in this specific metric. While Macrel recorded the highest overall accuracy (0.9908), AUC (0.9998), and specificity (1.0000), AMPBAN remains a highly reliable model with superior class balance compared to other established tools like AMPlify. AMPlify ranked third (Acc: 0.9413, MCC: 0.8885) but trailed AMPBAN by 3.1% in accuracy and 5.7% in MCC. These results underscore AMPBAN’s strong predictive power and its effectiveness on independently constructed data.

**Table 5.** Performance of AMPBAN compared with existing methods on XIAOAMP dataset. The bold values showed the best performance.

| Model | ACC | AUC | F1 | MCC | SN | SP |
| --- | --- | --- | --- | --- | --- | --- |
| CAMP3-ANN | 0.7995 | 0.7995 | 0.8161 | 0.6090 | 0.8902 | 0.7087 |
| AMPfun | 0.9168 | 0.9279 | 0.9229 | 0.8442 | 0.9957 | 0.8380 |
| AMPScanner | 0.7995 | 0.9361 | 0.8300 | 0.6419 | 0.9793 | 0.6196 |
| AI4AMP | 0.8484 | 0.9723 | 0.8658 | 0.7215 | 0.9783 | 0.7185 |
| AMPlify | 0.9413 | 0.9921 | 0.9445 | 0.8885 | **0.9989** | 0.8837 |
| Macrel | **0.9908** | **0.9998** | **0.9907** | **0.9817** | 0.9815 | **1.0000** |
| PTPAMP | 0.5495 | 0.7415 | 0.2231 | 0.1824 | 0.1293 | 0.9696 |
| smAMPsTK | 0.5491 | 0.7290 | 0.2058 | 0.2035 | 0.1164 | 0.9847 |
| AMP-CLIP | 0.8454 | 0.9299 | 0.8315 | 0.7121 | 0.9522 | 0.7740 |
| AMPBAN | 0.9723 | 0.9914 | 0.9729 | 0.9455 | 0.9946 | 0.9500 |

On the MFAAMP, AMPBAN achieved the highest accuracy (0.9525), MCC (0.9050), F1-score (0.9527), sensitivity (0.9496), and specificity (0.9554) (Table 6 and Figure 4D), outperforming all models, including strong contenders like AMPfun and AMPlify, on five of six metrics. The superior performance on a dataset with diverse peptide compositions further validates AMPBAN’s adaptability and precision.

**Table 6.**
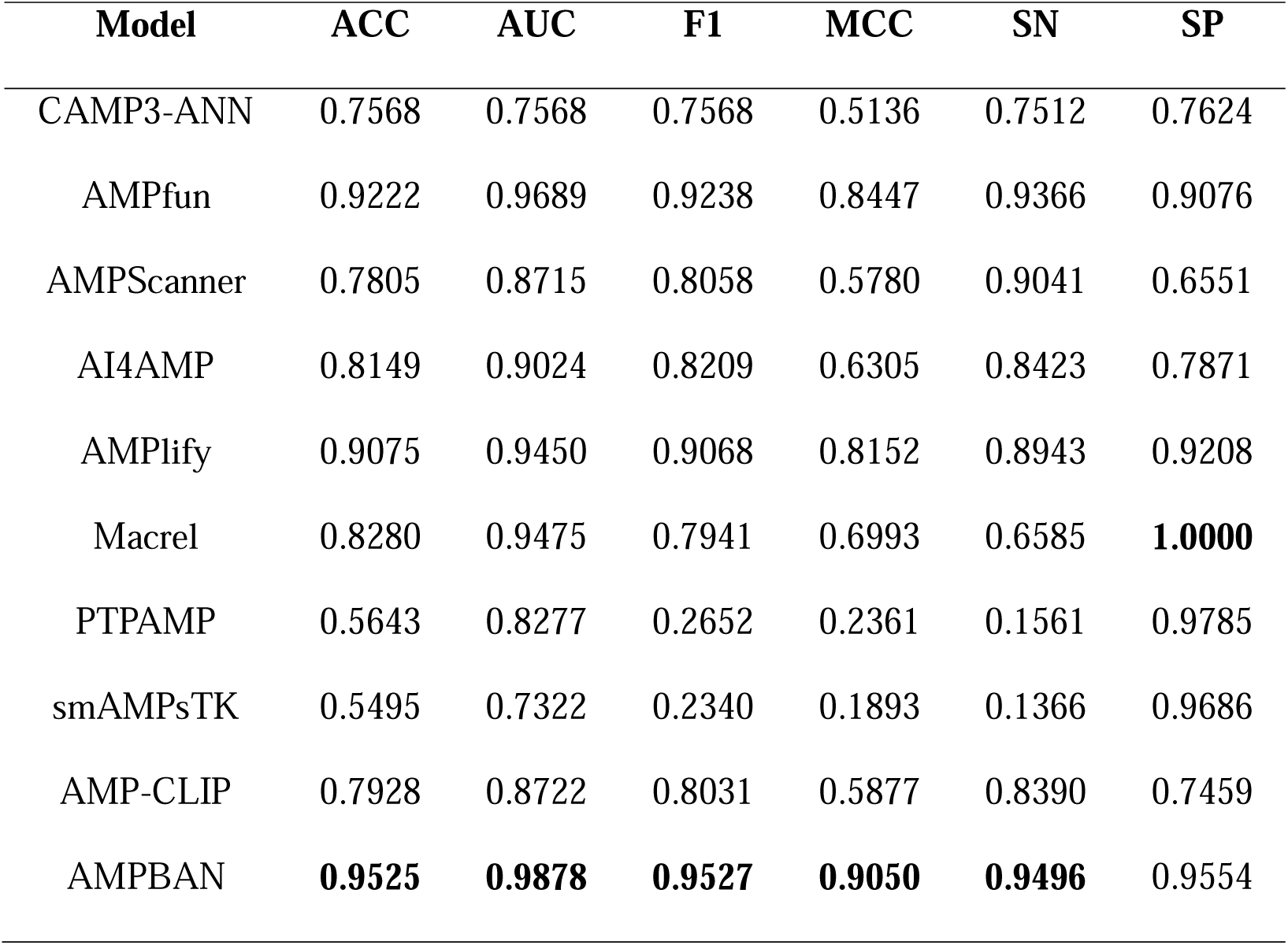
Performance of AMPBAN compared with existing methods on MFAAMP dataset. The bold values showed the best performance.

| Model | ACC | AUC | F1 | MCC | SN | SP |
| --- | --- | --- | --- | --- | --- | --- |
| CAMP3-ANN | 0.7568 | 0.7568 | 0.7568 | 0.5136 | 0.7512 | 0.7624 |
| AMPfun | 0.9222 | 0.9689 | 0.9238 | 0.8447 | 0.9366 | 0.9076 |
| AMPScanner | 0.7805 | 0.8715 | 0.8058 | 0.5780 | 0.9041 | 0.6551 |
| AI4AMP | 0.8149 | 0.9024 | 0.8209 | 0.6305 | 0.8423 | 0.7871 |
| AMPlify | 0.9075 | 0.9450 | 0.9068 | 0.8152 | 0.8943 | 0.9208 |
| Macrel | 0.8280 | 0.9475 | 0.7941 | 0.6993 | 0.6585 | **1.0000** |
| PTPAMP | 0.5643 | 0.8277 | 0.2652 | 0.2361 | 0.1561 | 0.9785 |
| smAMPsTK | 0.5495 | 0.7322 | 0.2340 | 0.1893 | 0.1366 | 0.9686 |
| AMP-CLIP | 0.7928 | 0.8722 | 0.8031 | 0.5877 | 0.8390 | 0.7459 |
| AMPBAN | **0.9525** | **0.9878** | **0.9527** | **0.9050** | **0.9496** | 0.9554 |

Overall, across all independent test datasets, AMPBAN demonstrated robust generalization and clear improvements over both traditional and deep learning–based predictors. These results confirm the effectiveness of incorporating evolutionary sequence features, structural representations, and bilinear attention mechanisms for AMP prediction.

### 3.5 Ablation Study

To systematically assess the contribution of each component in AMPBAN, we conducted an ablation study across four independent test datasets: Merged-AMP, MFAAMP, XIAOAMP, and XUAMP. We compared the full AMPBAN model against three variants: Sequence-only (using ESM3 sequence embeddings alone), Structure-only (using EGNN-derived structural features alone), and Sequence-Structure (concatenating both feature types before final classification without joint features). Six metrics were evaluated: Accuracy (ACC), AUC, F1, MCC, Sensitivity (SN), and Specificity (SP). ROC curves further highlight discriminative performance across datasets (Figure 5).

**Figure 5.**
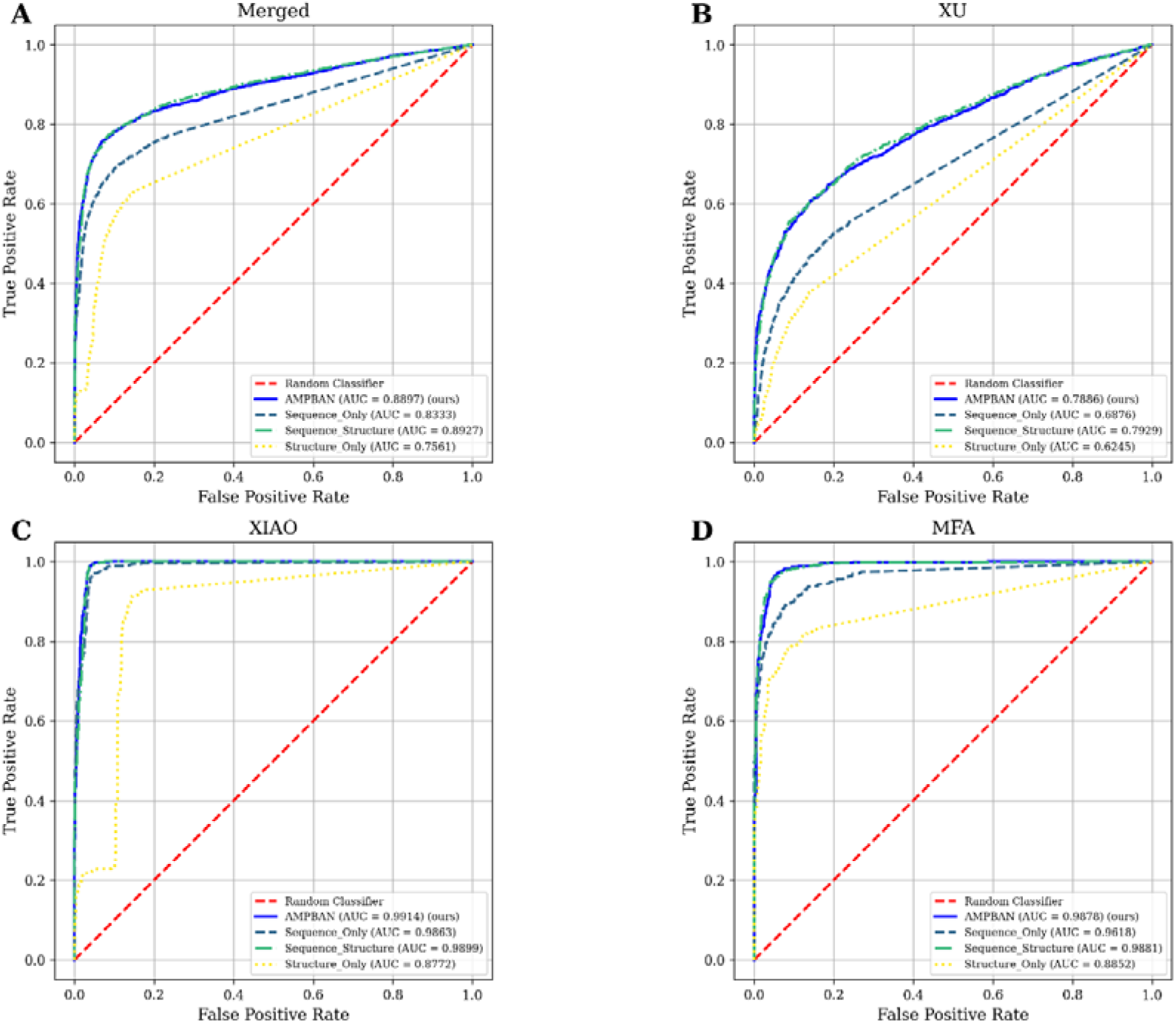
Receiver operating characteristic (ROC) curves of AMPBAN and the other three variants’ predictors on the four independent test datasets. (A) Merged-AMP: A combined dataset consisting of the XUAMP, XIAOAMP, and MFAAMP. (B) XUAMP. (C) XIAOAMP. (D) MFAAMP. The dotted red line represents the performance of a Random Classifier (AUC = 0.5); the blue line denotes AMPBAN.

The evaluation results revealed a distinct hierarchy in the performance of the models. Sequence-only achieved moderate accuracy (0.6579 on XUAMP to 0.9054 on XIAOAMP) with strong AUC on XIAOAMP (0.9863) but limited MCC (0.3227–0.8244) on harder datasets (Figure 5; Table 7). Structure-only was consistently weaker, with accuracy from 0.6175 (XUAMP) to 0.8745 (XIAOAMP) and MCC from 0.2658 to 0.7533, reflecting poor class balance (Figure 5; Table 7). It signified that single-modality models underperformed. The Sequence-Structure variant significantly improved performance through feature complementarity: ACC and AUC reached 0.6947 and 0.7929 on XUAMP, respectively; ACC and AUC were 0.9750 and 0.9899 AUC on XIAOAMP, respectively (Figure 5; Table 7). It demonstrated that combining sequence and structural signals markedly enhances prediction.

**Table 7.** Performance of AMPBAN compared with three variant models, *i.e.*, Sequence-only (ESMC-only), Structure-only (EGNN-only), and Sequence-Structure (ESMC-EGNN), across four independent test datasets. The bold values showed the best performance. ESMC-only: using sequence embeddings alone; EGNN-only: using structural features alone; ESMC-EGNN: combining features without BAN; AMPBAN: integrating sequence, structure, and BAN fusion.

| Dataset | Model | ACC | AUC | F1 | MCC | SN | SP |
| --- | --- | --- | --- | --- | --- | --- | --- |
| Merged-AMP | ESMC-only | 0.7686 | 0.8333 | 0.7694 | 0.5373 | **0.7708** | 0.7665 |
|  | EGNN-only | 0.7376 | 0.7561 | 0.7087 | 0.4854 | 0.6373 | 0.8383 |
|  | ESMC-EGNN | 0.8281 | **0.8927** | 0.8023 | 0.6807 | 0.6962 | **0.9605** |
|  | AMPBAN | **0.8358** | 0.8897 | **0.8157** | **0.6887** | 0.7258 | 0.9461 |
| XUAMP | ESMC-only | 0.6579 | 0.6876 | 0.6185 | 0.3227 | **0.5547** | 0.7611 |
|  | EGNN-only | 0.6175 | 0.6245 | 0.5011 | 0.2658 | 0.3841 | 0.8509 |
|  | ESMC-EGNN | 0.6947 | **0.7929** | 0.5864 | 0.4569 | 0.4329 | **0.9564** |
|  | AMPBAN | **0.7077** | 0.7886 | **0.6192** | **0.4691** | 0.4753 | 0.9401 |
| XIAOAMP | ESMC-only | 0.9054 | 0.9863 | 0.9133 | 0.8244 | 0.9957 | 0.8152 |
|  | EGNN-only | 0.8745 | 0.8772 | 0.8809 | 0.7533 | 0.9283 | 0.8207 |
|  | ESMC-EGNN | **0.9750** | 0.9899 | **0.9753** | **0.9503** | 0.9870 | **0.9630** |
|  | AMPBAN | 0.9723 | **0.9914** | 0.9729 | 0.9455 | **0.9946** | 0.9500 |
| MFAAMP | ESMC-only | 0.8411 | 0.9618 | 0.8606 | 0.7070 | **0.9740** | 0.7063 |
|  | EGNN-only | 0.8337 | 0.8852 | 0.8348 | 0.6675 | 0.8341 | 0.8333 |
|  | ESMC-EGNN | 0.9427 | **0.9881** | 0.9417 | 0.8864 | 0.9187 | **0.9670** |
|  | AMPBAN | **0.9525** | 0.9878 | **0.9527** | **0.9050** | 0.9496 | 0.9554 |

AMPBAN exhibited robust and balanced predictive capabilities across various datasets. On the Merged-AMP, AMPBAN achieved an ACC of 0.8358 and an MCC of 0.6887, surpassing the Sequence-Structure variant, which recorded an ACC of 0.8281 and an MCC of 0.6807. On the MFAAMP, AMPBAN reached an ACC of 0.9525 and an MCC of 0.9050, improving over Sequence-Structure’s ACC of 0.9427 and MCC of 0.8864. On XIAOAMP, AMPBAN delivered a high ACC of 0.9723, an AUC of 0.9914, and an MCC of 0.9455, nearly matching the Sequence-Structure accuracy but with a better balance (SP: 0.9500 vs. 0.9630). Even on the challenging XUAMP dataset, AMPBAN attained an ACC of 0.7077 and an MCC of 0.4691, substantially outperforming Sequence-Structure, which achieved an ACC of 0.6947 and an MCC of 0.4569, as well as the single-modality models.

These results confirm that sequence and structural representations capture complementary AMP characteristics, and the simple concatenation of both features already boosts performance. However, AMPBAN’s bilinear attention-based fusion further refines this integration, yielding superior accuracy, balance, and generalization, especially on diverse or difficult datasets. This ablation study validates the critical role of principled multi-modal fusion in achieving state-of-the-art AMP prediction.

## 4 Conclusion

This study developed a robust computational framework for AMP prediction by integrating sequence and structural features to address the limitations of sequence-centric models, accelerating AMP discovery in response to the global antimicrobial resistance crisis. The developed multi-modal model, AMPBAN, advances the field in two aspects: adopting ESM3 sequence embeddings with structural features derived from ESMFold and processed by Equivariant Graph Neural Networks, and employing a novel Bilinear Attention Network to model precise cross-modal interactions. The efficacy of AMPBAN was rigorously demonstrated by its superior and reproducible performance, consistently surpassing four re-implemented state-of-the-art models in 5-fold cross-validation, and achieving top-tier results against nine advanced predictors across diverse external test sets. This comprehensive design, underpinned by validated feature analysis, confirms the efficacy of our multi-modal fusion approach as a significant step forward in AMP prediction.

Despite its enhanced predictive strength, AMPBAN faces several practical limitations. The necessity of generating and processing 3D structures via ESMFold and EGNNs introduces a substantial increase in computational complexity, which may restrict its scalability for extremely large-scale virtual screening. Critically, the ablation study revealed lower sensitivity of AMPBAN in some cases (particularly in complex datasets like XUAMP), suggesting potential overfitting to specific characteristics—a challenge that may stem from the intricate integration of sequence and structural features. In fact, simpler models like the Sequence-Structure (ESMC-EGNN) variant sometimes delivered more balanced performance, indicating that lowering model complexity may improve robustness in certain cases. Finally, while highly effective at binary classification, the current model lacks the necessary granularity to distinguish between specific functional AMP classes (e.g., antibacterial vs. antifungal), limiting its direct utility for targeted drug design.

Looking ahead, future efforts could be diverted to optimize the model architecture to target for a better balance between complexity and generalization, thus addressing the potential overfitting issues observed in the ablation study. We plan to integrate additional structural descriptors, such as surface features [27] or solvent accessibility, and extend AMPBAN from a binary classifier to a multi-class prediction engine capable of forecasting specific functional activities. Concurrently, efforts will be directed toward computational optimization techniques to enhance efficiency, ultimately facilitating the clinical translation of AMP prediction and inspiring innovative strategies for peptide-based drug development.

### Key points

- We developed AMPBAN, a deep learning framework that seamlessly integrates evolutionary sequence features (from ESM3) and 3D structural features (encoded by Equivariant Graph Neural Networks, EGNNs) for accurate antimicrobial peptide prediction.
- AMPBAN utilizes a Bilinear Attention Network to explicitly model and fuse the complex, pairwise interactions between sequence and structural channels, generating a high-fidelity joint representation.
- Rigorous benchmarking and cross-validation demonstrated that AMPBAN consistently outperforms nine state-of-the-art predictors across diverse external test sets, underscoring its superior predictive capability and robustness.

## Author contributions

L.W. and L.B. conceived and designed the study. W.H.B. and W.H.Y. collected the data. W.H.B. and H.C.J. built the model and performed data analyses. W.H.B., W.H.Y., H.C.J., Y.Q.C., L.B. and L.W. wrote the manuscript. All authors read and approved of the final manuscript.

## Data availability

The data used in this study and source code are openly available for those interested at https://github.com/baiwenhuim/ampban.

