## Supplementary material for "AMPBAN: A Deep Learning Framework Integrating Protein Sequence and Structural Features for Antimicrobial Peptide Prediction": Supplementary Materials.docx

**Text S1:** Model implementation procedures.

The AMPBAN framework is implemented using PyTorch (v2.5.1) with Python 3.10. All computational experiments were performed on a workstation equipped with an NVIDIA GeForce RTX 3090 GPU (24 GB), running Ubuntu 24.04 via the Windows Subsystem for Linux (WSL2). To ensure reproducibility, a random seed of 42 was utilized for all operations, including data splitting and weight initialization. For sequence encoding, we utilized the ESMC model (Synthyra/ESMplusplus_small) to generate peptide representations. The structure embedding of peptides was addressed by EGNN module from a Python package ‘progres’ (<https://github.com/greener-group/progres>) [48]. The model architecture centers on a Bilinear Attention Network (BAN) that learns joint representations between the 128-dimensional structural features and 1920-dimensional sequence features. The BAN layer uses a hidden dimension of 128 and a bilinear value ($k$) of **3**. A multi-layer perceptron (MLP) with a hidden dimension of 128 and a Dropout rate of 0.4 processes the concatenated sequence, structure, and joint features. AMPBAN was trained to minimize the Binary Cross-Entropy (BCE) loss between predicted probabilities and ground-truth labels. Optimization was performed using the Adam optimizer with an initial learning rate of 1e-5 and an L2 weight decay of 1e-5 to encourage weight sparsity and prevent overfitting. Training was conducted with a batch size of 128 for a maximum of 200 epochs. We implemented an early stopping strategy that monitored validation loss with a patience of 5 epochs; the model state achieving the minimum validation loss was preserved for final evaluation.

**Text S2: Benchmark dataset filter criteria**

To ensure a high degree of confidence in their non-antimicrobial nature, we applied rigorous filtering criteria. This involved the exclusion of any sequences containing one or more of 16 predefined antimicrobial-related keywords, including terms such as "antimicrobial," "antibiotic," "antibacterial," "antiviral," "antifungal," "antimalarial," "antiparasitic," "anticancer," "defensin," "cathelicidin," "bacteriocin," "microbicidal," "fungicide," "excreted," and "effector." Furthermore, we eliminated duplicate sequences and those incorporating non-standard amino acids.

**Text S3: Calculate physical-chemical properties**

Physicochemical properties of the AMPs and NAMPs were computed using the ‘ProtParam.ProteinAnalysis’ class from Biopython. For each peptide sequence, the following properties were calculated: sequence length, net charge at pH 7.0, molecular weight, isoelectric point, instability index (Any value above 40 means the protein is unstable), Grand Average of Hydropathicity (GRAVY) according to Kyte and Doolitle, and secondary structure fractions (helix and sheet). The amino acid composition of AMPs and non-AMPs was determined by aggregating all sequences within each dataset into a single sequence and computing the percentage frequency of each amino acid using ‘ProtParam.ProteinAnalysis.get_amino_acids_percent’. This provided a comprehensive view of compositional differences between the two peptide classes.

To assess differences in physicochemical properties between AMPs and non-AMPs, the Mann-Whitney U test was applied using the ‘scipy.stats.mannwhitneyu’ function from SciPy (version 1.14.1). This non-parametric test was chosen to compare the distributions of each property (length, net charge, molecular weight, isoelectric point, instability index, GRAVY, helix fraction, and sheet fraction) between the two groups, accommodating potential non-normality in the data. The test was performed with a two-sided alternative hypothesis, and p-values were reported with significance levels denoted as follows: $p >0.05$ (not significant, ns), $p \leq0.05$ (*), $p \leq0.01$ (**), $p \leq0.001$ (***), and $p \leq0.0001$ (****).

**Text S4:** Detailed Implementation of Five-Fold Cross Validated Models.

To ensure the reproducibility and fairness of comparative experiments, we re-implemented the four SOTA models strictly following the original design principles and technical specifications described in its source publication, while adapting it to a standardized 5-fold cross-validation (CV) framework for unified evaluation with 6 metrics (sensitivity, specificity, accuracy, MCC, AUC, and F1-score) across all models. Fixed training data paths to the unified AMPBAN dataset. Below is the detailed implementation protocol, we made the following minimal, non-architectural modifications (Retained all original hyperparameters and sequence processing logic):

**AMPScanner (v2)**

1. Removed dependency on separate test/validation FASTA files (uses a single combined dataset for CV).
2. Added stratified 5-fold splitting (StratifiedKFold) with parameter (shuffle=True, random_state=123) to maintain class balance across folds.
3. Extended metric calculation to include Specificity.
4. Suppressed verbose training output (verbose=0) for cleaner CV results.

**AMP-CLIP**

1. Replaced CSV-based data loading with direct FASTA parsing.
2. Extended maximum sequence length from 100 to 200 AA (to match other models in the comparison; no impact on model architecture).
3. Added stratified 5-fold splitting (StratifiedKFold) to replace the fixed train/validation/test split in the original script.
4. Extended metric calculation to include standardized AUC.
5. Adjusted batch size to 50% of the training fold size (avoids memory issues with variable fold sizes).

**AMPlify**

1. Replaced the original 5-fold ensemble training (on the training set) with stratified 5-fold CV (train/validation split on the full dataset) to match evaluation protocols of other models.
2. Enforced CPU-only execution (via TensorFlow config) to eliminate stochasticity from GPU parallelism (critical for reproducibility).
3. Extended metric calculation to include MCC and standardized AUC.
4. Reduced patience from 50 to 5 (accelerates training without loss of performance; restores best weights to avoid overfitting).

**Macrel**

1. Replaced the fixed train/test split with stratified 5-fold CV (StratifiedKFold).
2. Extended metric calculation to include Specificity and AUC.
3. Adjusted RF random seeds (42+fold) to ensure independent tree initialization across folds.
4. Removed OOB prediction replacement (unnecessary for CV, as folds are non-overlapping) while retaining oob_score=True to match original RF configuration.
5. Saved one ONNX model per fold (instead of a single model) to enable traceability of CV results.
6. Explicitly encoded labels as 1 (AMP)/0 (non-AMP).

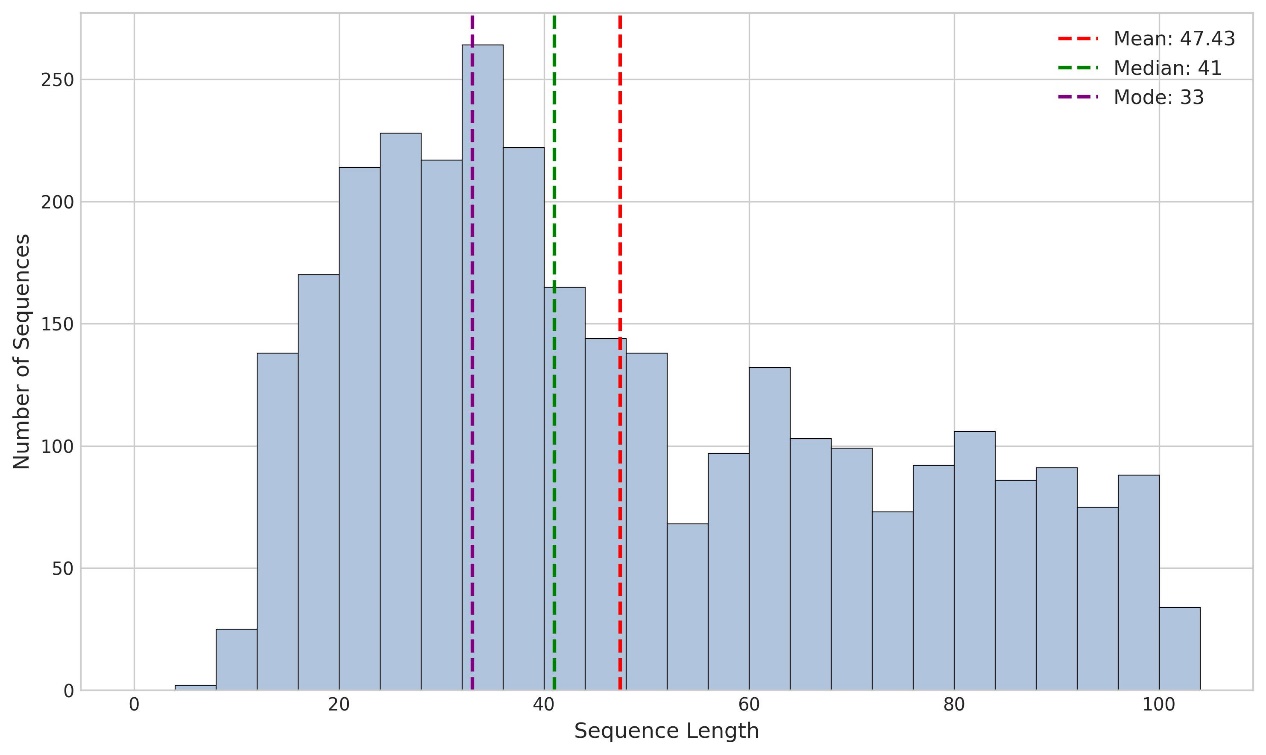

**Figure S1.** Distribution of sequence lengths for AMP sequences from Merged-AMP. Vertical dashed lines indicate key statistical metrics: the red line marks the mean sequence length (±0.01 residues), the green line indicates the median sequence length, and the purple line denotes the mode (most frequently observed sequence length).

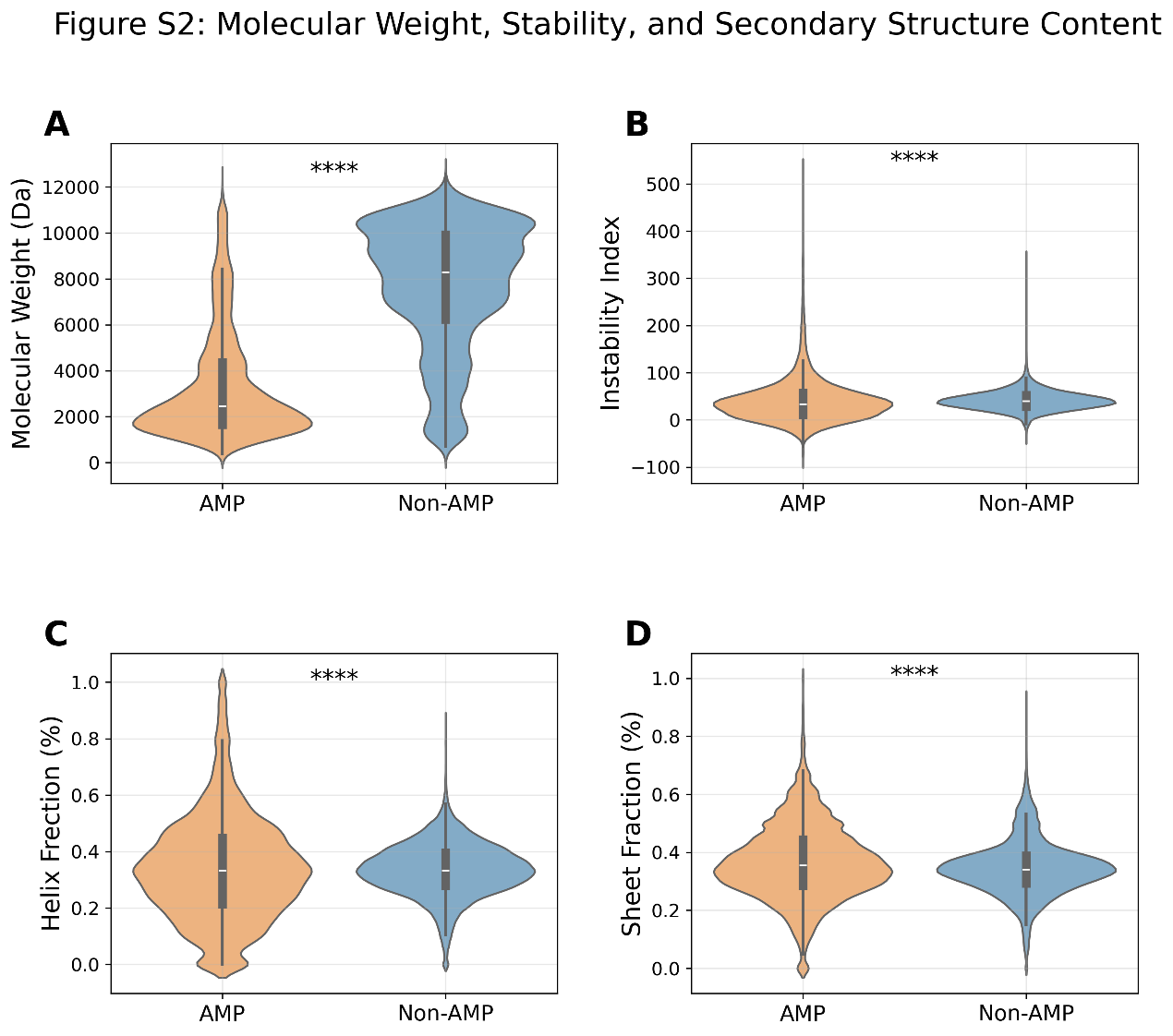

**Figure S2.** Distribution of physicochemical properties for AMPs and Non-AMPs. All panels display violin plots with internal boxplots (median = black line, interquartile range = gray bar); **** denotes *P* < 0.0001. (A) Molecular Weight (Da): The mass of the peptide/protein, measured in Daltons. (B) Instability Index: A metric predicting the in vitro half-life of a peptide. (C) Helix Fraction (%): The percentage of the peptide sequence that adopts an α-helical secondary structure. (D) Sheet Fraction (%): The percentage of the peptide sequence that adopts a β-sheet secondary structure.

**Table S1.** Summary of the nine state-of-the-art (SOTA) antimicrobial peptide prediction models used for performance comparison.

| Methods | Time | Model | Web Server or codes |
| --- | --- | --- | --- |
| CAMP3-ANN | 2015 | ANN | <http://www.camp3.bicnirrh.res.in/predict/> |
| AMPfun | 2020 | RF | <http://fdblab.csie.ncu.edu.tw/AMPfun/runClassifier.php> |
| AMPScanner | 2018 | CNN, LSTM | <https://www.dveltri.com/ascan/v2/ascan.html> |
| AI4AMP | 2021 | LSTM | <https://axp.iis.sinica.edu.tw/AI4AMP/> |
| AMPlify | 2022 | Bi-LSTM, Attention | https://github.com/bcgsc/AMPlify |
| Macrel | 2020 | RF | https://github.com/BigDataBiology/macrel |
| PTPAMP | 2023 | SVM | <http://14.139.61.8/PTPAMP/prediction.php> |
| smAMPsTK | 2024 | SVM | https://github.com/skbinfo/smAMPsTK |
| AMP-CLIP | 2025 | CNN, LSTM, Attention, Transformer | https://github.com/MicroResearchLab/AMP-potency-prediction-EvoGradient |
